# Protein Predictors of Worsening Cerebral Edema in Traumatic Brain Injury in Critically Ill Patients

**DOI:** 10.64898/2026.05.23.727416

**Authors:** Hannah L. Radabaugh, Ahmed Abdelhak, Kiarra Ning, Ruchira M. Jha, Susan E. Rowell, Jeffrey M. Pollock, Esmeralda Mendoza, Luisa M. Rojas Valencia, Adam R. Ferguson, HE Hinson

## Abstract

Cerebral edema (CE) is a major determinant of poor outcomes after traumatic brain injury (TBI). CE affects approximately 60% of patients with mass lesions on head computed tomography (hCT) and confers a tenfold increase in mortality. Despite this burden, no validated biomarkers exist to identify patients at risk for early CE worsening or to elucidate the biological processes underlying divergent edema trajectories. We conducted high-throughput plasma proteomic profiling in a prospective moderate–severe TBI cohort with evidence of intracranial blood on the initial post-injury hCT scan (n=123 patients) to (1) identify a clinically feasible set of biomarkers plus a predictive model that could be used to assess CE risk at admission, and (2) conduct a mechanistic, hypothesis-generating analysis to identify the molecular networks involved in CE progression in the first hours post-injury. Plasma collected within 3 hours of injury was analyzed using the Olink Explore platform (5394 proteins), and patients underwent head CT imaging upon presentation and at 6 hours. Two complementary analytic pathways, reflecting the study’s two goals, were pursued in parallel. First, logistic regression with false discovery rate (FDR) correction identified a conservative 12-protein set associated with baseline CE. When combined with targeted clinical variables (i.e., age, sex, admission GCS, pupil reactivity, admission glucose, admission alcohol), this panel supported high-performing supervised classifiers for predicting CE worsening by 6 hours post-baseline scan (best model: XGBoost, AUC = 0.78; recall = 0.83). Second, a broader 60-protein panel, selected via bootstrapped elastic-net regression, was used to interrogate the mechanistic architecture of CE worsening using random forest SHAP attribution and protein–protein interaction modeling. Proteomic signatures diverged sharply between patients whose CE did and did not worsen. The CE worsening group was characterized by a coherent neuronal–synaptic injury axis dominated by ELAVL4, CEND1, NEFL, NECAB2, GFAP, CHGB, RPH3A, HPCAL4, and HOMER1, proteins involved in neuronal structural integrity, vesicular trafficking, synaptic vesicle cycling, calcium signaling, and axonal degeneration. These reciprocal proteomic states suggest that early edema progression may be driven by coordinated disruption of neuronal and synaptic resilience programs. Together, these findings suggest (1) a hyperacute biomarker panel plus predictive model with potential for prospective validation and (2) mechanistic evidence for a distinct neuronal–synaptic injury network that associates with early edema worsening after TBI.

## Introduction

Traumatic brain injury (TBI) initiates a rapidly evolving cascade of secondary injury processes that shape early mortality and long-term neurological disability. Among these processes, brain swelling or cerebral edema (CE) is one of the most consequential, contributing directly to acute neurological decline.^1–3^ Despite its central role in early deterioration, CE remains challenging to predict and monitor. Current clinical practice relies almost entirely on serial neurological exams, serial imaging, and in severe cases invasive ICP monitoring.^4,5^ These methods are imperfect as they only allow clinicians to detect worsening edema after it is well established, and in the case of imaging, after structural changes have become radiographically apparent. By this stage, the biological drivers of progression may have intensified, narrowing the therapeutic window and limiting opportunities for early intervention.

The absence of validated time-of-admission biomarkers capable of predicting early CE worsening reflects a broader limitation in neurotrauma management. Although TBI is increasingly understood as a systems-level disorder involving complex interactions among neuronal, glial, vascular, and immune processes, clinical monitoring still depends heavily on gross structural changes rather than on underlying biology. Numerous studies have implicated ionic imbalance, cytoskeletal breakdown, astrocytic reactivity, excitotoxicity, endothelial dysfunction, oxidative stress, and inflammation as contributors to CE, yet these pathways have rarely been assessed in combination or within the hyperacute clinical window in which patients diverge toward worsening or stability.^3,6–8^ Consequently, clinicians lack tools to identify which patients are likely to deteriorate in the acute phase, limiting timely decisions regarding monitoring intensity, neurosurgical readiness, or early therapeutic strategies.

Advances in high-throughput proteomics now offer an opportunity to overcome decision support barriers. Modern platforms such as Olink Explore proximity extension assays allow simultaneous measurement of thousands of circulating proteins with high sensitivity and reproducibility, opening new avenues for both clinical biomarker discovery and mechanistic investigation.^9–11^ Plasma proteomics is particularly well suited to acute TBI, as it enables minimally invasive characterization of molecular injury states within hours of trauma, well before radiographic changes emerge and without the need for patient transport to a scanner. When integrated with machine learning (ML), proteomic datasets can support not only the identification of clinically actionable protein signatures but also the mapping of biological networks that may differentiate dynamic edema trajectories. Yet, despite the clear potential of this approach, large-scale proteomic studies focused on well characterized TBI cohorts remain limited. In the current work we applied a modern 5400 protein assay to further development of plasma biomarkers in GCS 3-12 TBI patients (previously known as ‘moderate’ to ‘severe’).

To address these gaps, we performed high-throughput plasma proteomic profiling using samples from the previously conducted prospective, observational <u>PR</u>edictors <u>O</u>f low-risk phenotypes after <u>T</u>raumatic brain injury <u>I</u>ncorporating <u>P</u>roteomic biomarker <u>S</u>ignatures (PROTIPS study).^12^ The cohort for the current analysis consisted of adults (ages 17-85) with CT-positive (intra- or extra-axial hemorrhage on the initial post-injury head CT), moderate–severe (GCS 3-12) TBI who underwent standardized CT imaging at admission and a per-protocol follow-up CT 6 hours later to examine for hemorrhage expansion. The analytical framework developed for this post-hoc, hypothesis-generating study was structured around two complementary aims designed to bridge clinical utility and mechanistic biology. The first goal focused on clinical decision support: to identify a clinically feasible biomarker panel and develop a predictive model with plausible translational potential for admission-time CE risk assessment. Such a tool could ultimately guide escalation of monitoring, inform early treatment decisions, and provide a biologically grounded alternative to dependence on episodic CT imaging. The second goal was mechanistic and hypothesis-generating: to use a broader proteomic signal to explore the molecular architecture underlying divergent edema trajectories and to identify coordinated biological networks that may drive progression in the hyperacute post-injury period.

In pursuing these aims, we leveraged modern statistical and ML approaches capable of handling high-dimensional biological data, stabilizing feature selection, and quantifying directional contributions of proteins to model predictions. This dual analytic framework was chosen deliberately: clinical translation requires parsimony, feasibility, and interpretability, whereas mechanistic discovery benefits from broader feature space exploration and network-level interrogation. In sum, the biological complexity and clinical urgency of CE demand approaches that integrate deep molecular profiling with predictive modeling. Developing tools capable of anticipating CE worsening and identifying the biological programs that may underlie this progression, represent critical steps toward more proactive and biologically informed management of acute TBI.

## Materials and Methods

### Participants

This study is a post-hoc analysis of the previously collected PROTIPS cohort. PROTIPS was a prospectively enrolled, single center, observational cohort study of adults and older teenagers (age>14) suffering moderate to severe (GCS 3-12) TBI and intracranial hemorrhage on the initial head CT. Trauma research coordinators with formal GCS training screened sequential patients up to 3 hours from injury for inclusion. Exclusion criteria consisted of GCS score of 3 without a reactive pupil, unknown time of injury, prisoners, pregnant individuals, or no hemorrhage on Head CT. Consent for participation was obtained from participant’s legally authorized representative. If/when the participant regained the ability to provide informed consent, consent was re-obtained. Demographics, injury characteristics, and hospital course were abstracted from the electronic health record by trained research coordinators. Blood was drawn at the time of admission (within 3 hours of injury), centrifuged, aliquoted for plasma, and immediately stored at -80°C. Institution review board (IRB) approval was obtained prior to recruitment of participants.

### Imaging, Determination of Cerebral Edema (CE)

The primary outcome for analysis was the radiographic worsening of CE on a follow-up hCT performed 6 hours (average 5.62 hours) after the admission hCT, capturing the hyperacute phase of CE development.^6,13^ All hCTs (admission and 6 hour follow-up) were reviewed by a neuroradiologist for the presence CE. CE was assessed by examining the relevant components of the Marshall score basal cistern compression (absent/compressed/obliterated), midline shift (<5MM or >5mm).^14^ If the basal cisterns were compressed or obliterated, and/or midline shift was >5mm, the scan was judged to show cerebral edema (“CE present”). If CE worsened from the initial scan to the follow up scan, based on a progression between one or both of these categories (e.g. basal cistern compression increased from absent to compressed, or midline shift increased by any amount), the second scan was judged to show “worsening CE”. Of the 123 patients with complete proteomic data, 116 had a follow-up scan at 6 hours. Data missingness checks revealed this to be the only source of missingness in dataset.

### Plasma Proteomic Analysis

High-throughput proteomic analysis was performed using the Olink Explore HT platform, which utilizes proximity extension assay (PEA) technology to simultaneously measure approximately 5400 protein targets.^9,10^ All samples were processed according to standard manufacturer’s protocol. Protein abundances were reported as normalized protein expression (NPX) values generated using the Olink recommended data processing pipeline which incorporates internal and external assay control, across-plate normalization, and log2 transformation to yield relative protein abundance values. All 126 samples were processed within a single analytical batch. No samples failed Olink’s built-in quality control metrics (**Supplementary Fig. 1A**). As additional quality assessments, we examined the distribution of NPX values across samples, stratified by hemolysis index, and observed no systematic shifts in global protein abundance associated with hemolysis status (**Supplementary Fig. 1B**). Sample-level NPX distributions were highly comparable across all samples passing quality control. To further evaluate residual technical structure, we performed principal component analysis (PCA) on the NPX matrix. PCA revealed no grouping by quality control status and no dominant axes consistent with batch or technical artifacts, indicating that Olink’s standard processing adequately controlled for technical variation across samples (**Supplementary Fig. 1C).**

### Goal 1: Clinical-decision Making Support Biomarker Panel Screen

Goal 1 focused on deriving a clinically actionable biomarker panel for early risk stratification of CE worsening. To prioritize feasibility for potential downstream clinical translation and avoid overfitting, a conservative, hypothesis-agnostic discovery strategy was employed (**Fig. 1**; Modeling Phase). Univariate logistic regression was performed for each of the 5,394 measured proteins, using the presence of CE on admission head CT (present vs absent) as the binary outcome. To control for multiple testing and limit false discoveries, p-values were adjusted using the Benjamini–Hochberg false discovery rate (FDR) procedure, with a significance threshold of q < 0.05. Proteins meeting this criterion were retained, yielding a 12-protein panel associated with CE status at presentation.

**Figure 1.**
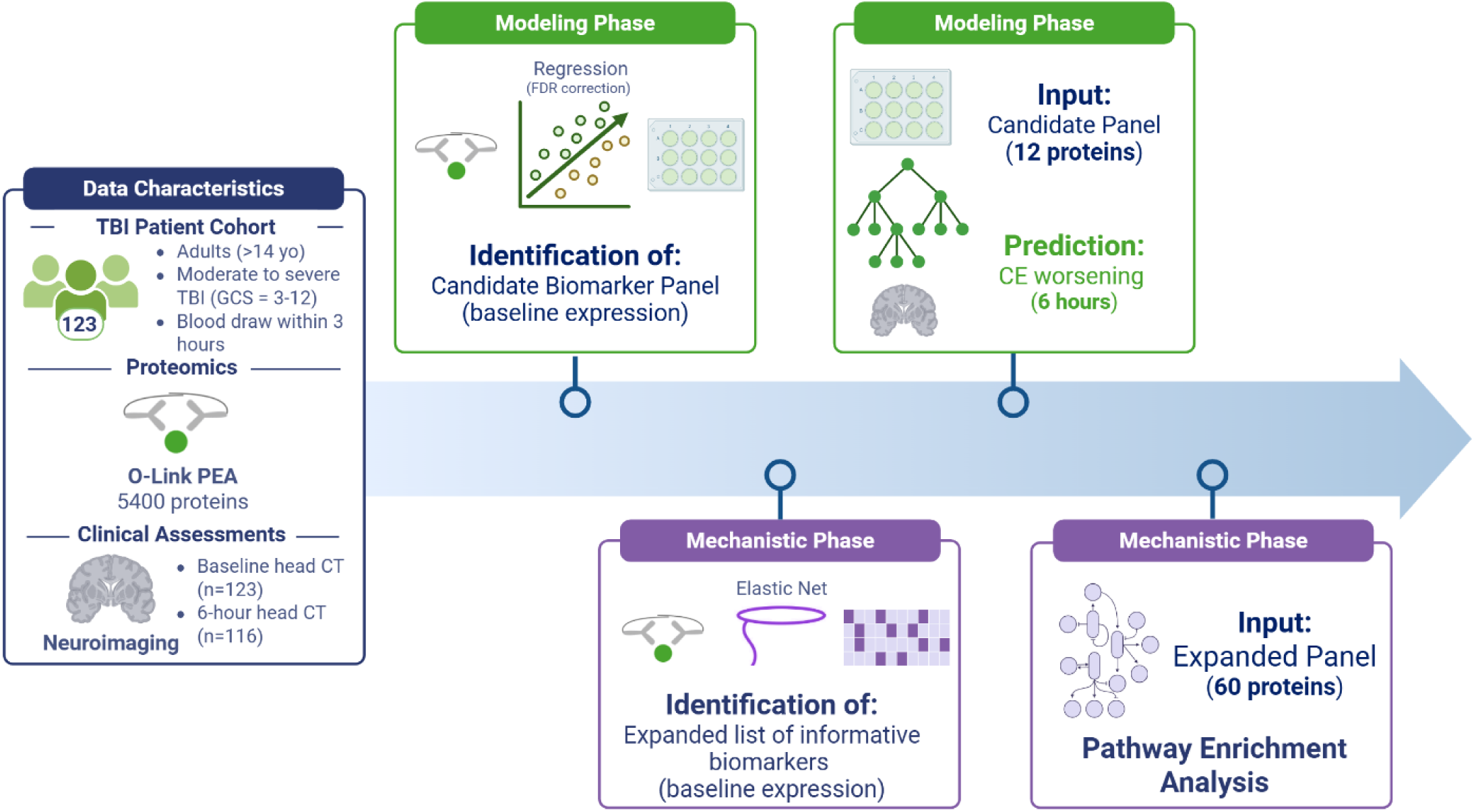
Study design and analytic workflow. Overview of clinical-decision making support (Goal 1; Green) and mechanistic (Goal 2; Purple) analyses integrating admission plasma proteomics and clinical data in patients with TBI. In Aim 1, univariate regression with FDR correction identified a parsimonious 12-protein panel associated with CE on admission CT, which was combined with admission clinical variables and evaluated using supervised ML to predict CE worsening at 6 hours. In Aim 2, bootstrapped elastic-net regression identified an expanded 60-protein panel for unsupervised structure analysis, model interpretation, and pathway enrichment.

### Supervised Classification

For predictive modeling of cerebral edema (CE) worsening at 6 hours, the 12-protein panel was integrated with six routinely available clinical variables obtained at hospital admission: age, sex, admission Glasgow Coma Scale (GCS) score, pupil reactivity, admission serum glucose, and admission blood alcohol level. These variables represent the final clinical feature set retained for modeling after evaluation of data completeness and stability during internal model development which included recursive feature selection with cross validation (RFECV). RFECV was tuned using the AUC scoring metric to determine the optimal set where adding more features no longer provide additional predictive value. The resulting combined feature set was used to capture complementary molecular and clinical information relevant to early pathophysiology and injury severity and to define a parsimonious input set suitable for external validation.

Four supervised machine-learning classifiers were trained and compared to identify the optimal predictive model: (1) penalized logistic regression with L2 (ridge) regularization, a generalized linear model that reduces overfitting by shrinking coefficient estimates; (2) random forest (RF), an ensemble of decision trees trained on bootstrapped samples that captures non-linear relationships and feature interactions; (3) support vector machine (SVM) with a radial basis function kernel, which constructs a non-linear decision boundary in a high-dimensional feature space; and (4) extreme gradient boosting (XGBoost), a gradient-boosted tree ensemble that iteratively improves predictive performance by correcting prior model errors.^15–19^ Model performance across classifiers was evaluated using the same feature inputs to facilitate consistent comparison and streamlined reporting.

### Model Performance

All models were trained on the combined clinical–proteomic feature set to predict binary radiographic CE at the 6-hour follow-up CT. Model development, internal validation, and performance reporting were conducted in accordance with established guidelines for transparent reporting of multivariable prediction models (TRIPOD and TRIPOD-AI).^20^ Internal validation was performed using a repeated, stratified, nested 5-fold cross-validation framework, with inner-loop cross-validation used for hyperparameter tuning and outer-loop cross-validation used to estimate out-of-sample performance, thereby minimizing information leakage and optimism bias.

Model performance was evaluated using multiple complementary metrics appropriate for binary classification, including the area under the receiver operating characteristic curve (AUC-ROC) to summarize discrimination across decision thresholds, the area under the precision–recall curve (PR-AUC) to account for outcome imbalance, and threshold-dependent measures of accuracy, sensitivity (recall), specificity, positive predictive value, and negative predictive value.^15,16^ The model selected for primary reporting was chosen based on overall discrimination, sensitivity for early clinical risk stratification, and stability of performance across cross-validation repeats. Differences in AUCs between correlated ROC curves were assessed using DeLong’s test, a nonparametric method for comparing model discrimination on the same set of subjects.

### Goal 2: Mechanistic Biomarker Panel Selection

To identify a broader proteomic signature associated with CE status and to support downstream mechanistic analyses at the individual patient level, we employed a feature-selection strategy distinct from Aim 1. Bootstrapped elastic-net logistic regression was applied with baseline CE presence on admission CT (present vs absent) as the binary outcome. Elastic net is a regularized regression approach that combines L1 (lasso) and L2 (ridge) penalties, enabling simultaneous variable selection and coefficient shrinkage while accommodating correlated predictors, a common characteristic of high-dimensional proteomic data.^21^

Models were fit across 1,000 bootstrap resamples using a mixing parameter of α = 0.5 to balance L1 and L2 regularization effects. Proteins selected in more than 70% of bootstrap iterations were retained. This panel was interpreted as representing core proteomic correlates of the acute CE state immediately post-injury.

### Modeling and Interpretation

Analyses proceeded in three stages to characterize the relationship between the 60-protein panel and subsequent CE worsening (**Fig. 1**; Mechanistic Phase). First, principal component analysis (PCA) was applied to the standardized protein abundances to perform unsupervised dimensionality reduction, summarizing correlated proteins into orthogonal principal components (PCs) that capture the dominant sources of variance in the data.^22^ Hierarchical clustering was then performed on protein co-expression patterns to identify groups of proteins exhibiting similar expression profiles, providing an unsupervised assessment of proteomic structure independent of outcome labels.

Second, supervised predictive modeling was used to test whether this baseline CE-associated proteomic signature could discriminate future clinical trajectories. The same suite of supervised ML classifiers was trained using only the 60-protein panel as input features to predict binary CE worsening at 6 hours post-injury. The best-performing model, as defined in Aim 1, was selected for downstream interpretation.

Model interpretation was performed using SHapley Additive exPlanations (SHAP), a framework derived from cooperative game theory that quantifies the marginal contribution of each feature to an individual prediction.^23^ SHAP values provide both the direction and magnitude of each protein’s contribution to predicted CE worsening risk on a per-subject basis, enabling consistent comparison of feature importance across samples.

Finally, proteins identified as high-impact contributors based on SHAP analyses were used as inputs for pathway enrichment and functional network annotation analyses.^24–27^ This approach enabled biological contextualization of model-derived signals by identifying overrepresented pathways and processes associated with CE pathophysiology.

## Results

### Demographics

Of the 123 patients with moderate-to-severe TBI and intracranial hemorrhage, 38 presented with CE on the baseline scan and 31% experienced radiographic worsening of CE by the 6-hour follow-up CT. Key demographic and clinical characteristics are presented in **Table 1**. The cohort had a median age of 43, was predominantly male, and had a median admission GCS score of 8. The groups with and without CE worsening were comparable in terms of established clinical risk factors (**Table 1**).

**Table 1:** Characteristics of patients, stratified by presence of CE on admission CT. Demographic and clinical characteristics of PROTIPS cohort. Continuous variables are presented as mean ± SD for normally distributed variables (Shapiro–Wilk p ≥ 0.05 in both groups) and as median [IQR] otherwise; compared using Student’s t-test or Mann–Whitney U test, respectively. Categorical variables are presented as n (%) and compared using χ2 test, or Fisher’s exact test where appropriate. †Follow-up scan was available for 116 of the 123 patients with Olink data available. ISS, Injury Severity Score; AIS, Abbreviated Injury Score; CT, computed tomography; GCS, Glasgow Coma Scale; INR, international normalised ratio; IQR, interquartile range; SDH, subdural hematoma. *Race “Other” comprises Asian (n=5), other (n=4), declined to answer (n=4), Native Hawaiian/Pacific Islander (n=2), and American Indian/Alaska Native (n=2). Ethnicity “Decline to answer” (n=3) excluded from ethnicity comparison.

|  | CE absent (n=85) | CE present (n=38) | *p* value |
| --- | --- | --- | --- |
| Age (years) | 42 [27–65] | 43 [25–56] | 0.359 |
| Sex |  |  | 0.120 |
| Male | 56 (65.9%) | 31 (81.6%) |  |
| Female | 29 (34.1%) | 7 (18.4%) |  |
| Race |  |  | 0.738 |
| White | 65 (76.5%) | 30 (78.9%) |  |
| Black | 7 (8.2%) | 4 (10.5%) |  |
| Other* | 13 (15.3%) | 4 (10.5%) |  |
| Ethnicity |  |  | 0.132 |
| Non-Hispanic | 79 (95.2%) | 32 (86.5%) |  |
| Hispanic | 4 (4.8%) | 5 (13.5%) |  |
| Admission GCS | 8 [7–10] | 7 [3–8] | 0.008 |
| GCS motor | 5 [4–6] | 4 [1–5] | 0.005 |
| ISS | 22 [14–29] | 26 [18–33] | 0.053 |
| AIS | 4 [3–4] | 4 [4–5] | <0.001 |
| Pupils |  |  | 0.079 |
| Both reactive | 71 (83.5%) | 27 (71.1%) |  |
| One reactive | 12 (14.1%) | 6 (15.8%) |  |
| Neither reactive | 2 (2.4%) | 5 (13.2%) |  |
| Hypotension | 13 (15.3%) | 4 (10.5%) | 0.671 |
| Hypoxia | 8 (9.4%) | 2 (5.3%) | 0.722 |
| Blood glucose (mg/dL) | 136 [115–166] | 135 [119–172] | 0.540 |
| Hemoglobin (g/dL) | 13.2 ± 1.7 | 12.9 ± 2.0 | 0.323 |
| Alcohol level (mg/dL) | 0 [0–195] | 0 [0–147] | 0.822 |
| Platelets (×10<sup>3</sup>/μL) | 242.5 ± 74.3 | 249.2 ± 69.9 | 0.632 |
| INR | 1.06 [1.01–1.17] | 1.07 [0.99–1.19] | 0.627 |
| Fibrinogen (mg/dL) | 282 [227–318] | 249 [186–290] | 0.042 |
| Craniotomy | 11 (12.9%) | 17 (44.7%) | <0.001 |
| In-hospital mortality | 7 (8.2%) | 15 (39.5%) | <0.001 |
| *Baseline CT findings* |  |  |  |
| SDH | 60 (70.6%) | 36 (94.7%) | 0.006 |
| Skull fracture | 39 (45.9%) | 34 (89.5%) | <0.001 |
| Basal cisterns |  |  | <0.001 |
| Normal | 75 (88.2%) | 19 (50.0%) |  |
| Compressed | 9 (10.6%) | 14 (36.8%) |  |
| Obliterated | 1 (1.2%) | 5 (13.2%) |  |
| Midline shift >5 mm | 7 (8.2%) | 18 (47.4%) | <0.001 |
| Largest hematoma volume, mL | 1.0 [0.0–3.0] | 3.0 [1.0–13.0] | <0.001 |
| Rotterdam score | 3 [3–3] | 4 [3–5] | <0.001 |
| *Follow-up CT (n=116)<sup>†</sup>* |  |  |  |
| Worsening edema on follow-up CT<sup>†</sup> | 13/79 (16.5%) | 23/37 (62.2%) | <0.001 |
| In-hospital mortality | 7 (8.2%) | 15 (39.5%) | <0.001 |

### A Parsimonious Admission-Time Biomarker Panel

The development of clinically useful candidate biomarkers requires a protein signature that is both biologically informative and feasible for measurement in the hyperacute setting. To this end, we first identified a concise plasma biomarker panel strongly associated with CE on the admission CT scan. Using a conservative logistic regression screen with FDR correction, we identified a 12-protein panel that robustly discriminated patients with and without CE at presentation (Aim 1).

The resulting panel was comprised of N-terminal EF-hand calcium binding protein 2 (NECAB2), hippocalcin-like protein 4 (HPCAL4), glial fibrillary acidic protein (GFAP), cell cycle exit and neuronal differentiation protein 1 (CEND1), ELAV-like RNA binding protein 4 (ELAVL4), glutamate decarboxylase 2 (GAD2), Homer scaffold protein 1 (HOMER1), N-terminal EF-hand calcium binding protein 1 (NECAB1), neurofilament light chain (NEFL/NFL), rabphilin 3A (RPH3A), glutamate decarboxylase 1 (GAD1), and synaptosome associated protein 25 (SNAP25) (**Fig. 2**).

**Figure 2.**
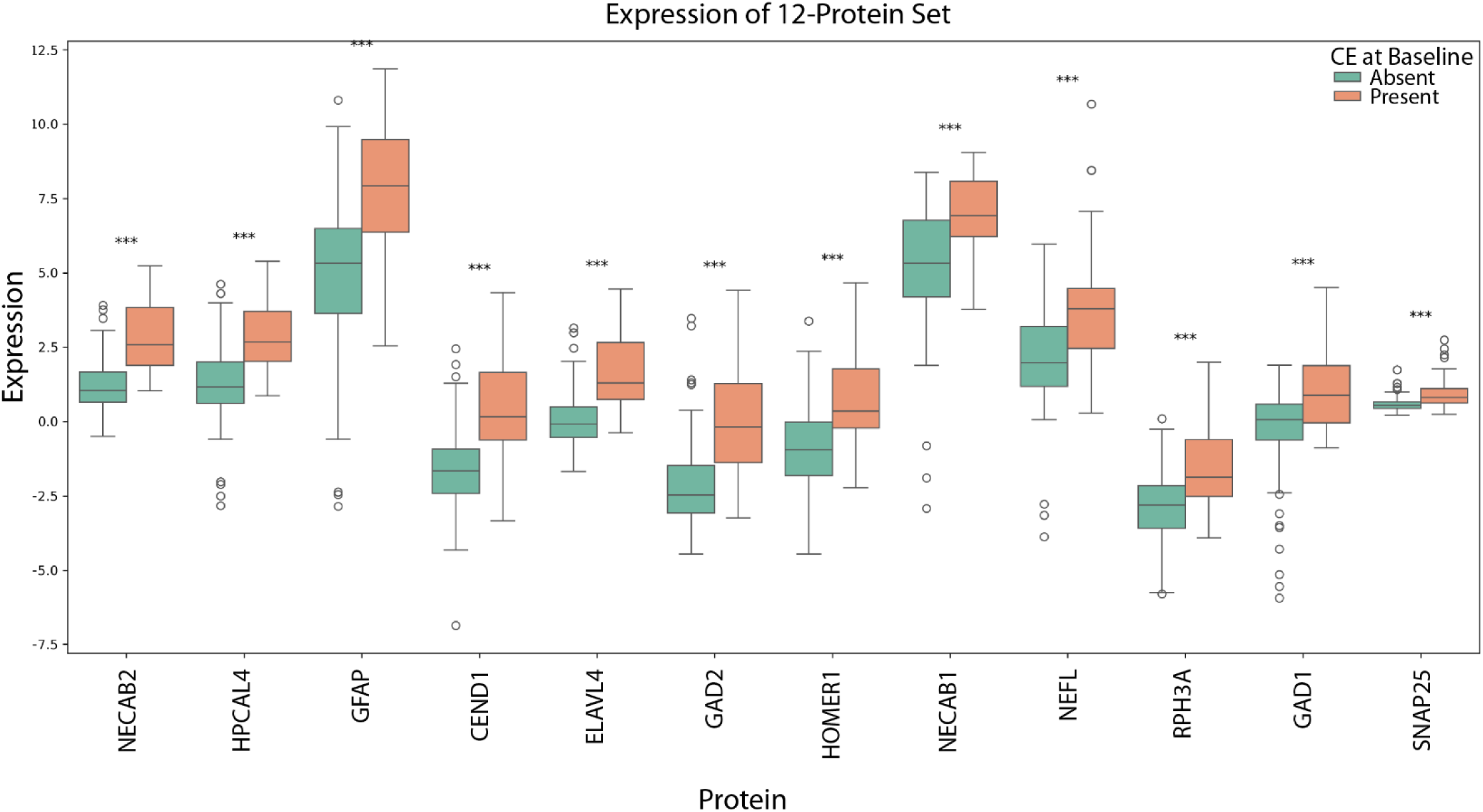
Differential expression of admission-time plasma proteins associated with cerebral edema. Boxplots show normalized plasma expression levels of the 12 proteins comprising the admission-time biomarker panel, stratified by presence or absence of CE on the admission CT scan. Boxes indicate the interquartile range with the median shown as a horizontal line; whiskers extend to 1.5x the interquartile range (IQR), and points denote outliers. Proteins were identified through univariate logistic regression with false discovery rate correction (q < 0.05). Asterisks denote statistically significant differences between CE-present and CE-absent groups (***q < 0.001).

Across the panel, protein expression levels differed significantly between patients with and without CE at admission, with several markers exhibiting marked shifts in central tendency and dispersion (**Fig. 2**). Notably, all proteins demonstrated coordinated alterations, suggesting that CE at presentation is accompanied by widespread disruption across multiple cellular compartments rather than isolated injury signals. This 12-protein panel was carried forward as the candidate molecular feature set for downstream predictive modeling in Aim 1 and evaluation of early CE worsening risk.

### A Predictive Model for Early CE Worsening

Recognizing that clinical decision-making integrates multiple data streams, we evaluated whether admission-time proteomic information improved prediction beyond routinely collected clinical variables. For predictive modeling of CE worsening, the 12-protein panel was combined with a parsimonious set of six admission clinical variables (age, biological sex, admission GCS score, pupil reactivity, admission serum glucose, and admission blood alcohol level) which represented the final clinical feature set retained following internal model development and stability assessment. This combined feature set was used consistently across models to facilitate interpretable comparison and to define clinically feasible input set for downstream validation.

Multiple supervised ML classifiers were trained to integrate clinical and proteomic features into a single risk estimate for radiographic CE worsening within 6 hours. Across modeling approaches, XGBoost demonstrated the most favorable balance of discrimination, recall, and stability and was therefore selected for primary reporting (**Fig. 3 A**). Using the combined clinical–proteomic feature set, the XGBoost model achieved good overall discrimination (AUC = 0.78, 95% CI [0.67-0.87]) and high sensitivity for early detection (recall = 0.83), correctly identifying 83% of patients who subsequently exhibited radiographic worsening. Importantly, incorporation of the proteomic panel provided a statistically significant improvement in predictive performance compared with a model using clinical variables alone (clinical-only AUC = 0.62; DeLong’s test, p < 0.01; **Fig. 3 B**). These results indicate that admission plasma protein measurements capture prognostic information not fully reflected in standard clinical assessments and meaningfully enhance early risk stratification for CE worsening.

**Figure 3.**
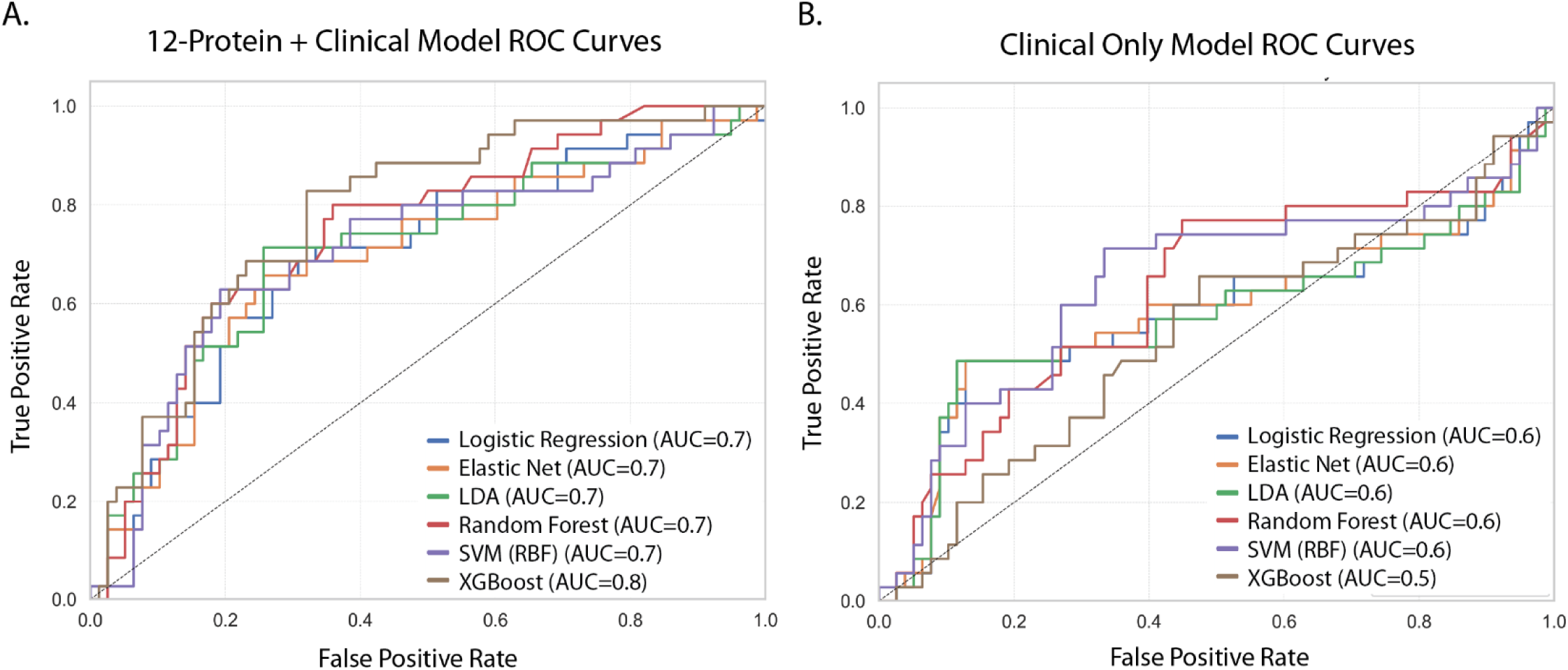
ROC curves comparing clinical–proteomic and clinical-only models for early CE worsening. ROC curves depicting model discrimination for prediction of radiographic CE worsening within 6 hours of admission head CT. **(A**) Models trained using the combined 12-protein admission-time biomarker panel and clinical variables demonstrate improved discrimination across multiple supervised learning algorithms, with XGBoost achieving the highest AUC (0.8). (**B**) Corresponding ROC curves for models trained using clinical variables alone show consistently lower discrimination across classifiers. Curves represent cross-validated model performance, with AUC values summarized in the legend. The dashed diagonal line indicates performance no better than chance.

### A Broad Proteomic Signature Associated with CE at Admission

To characterize the molecular architecture associated with CE on baseline head CT, we applied a complementary, multivariate feature-selection approach to identify a broader proteomic signature beyond the parsimonious panel derived in Aim 1. Using bootstrapped elastic-net regression with baseline CE presence as the outcome, we identified a stable panel of 60 proteins that were consistently selected across bootstrap iterations (>70%) and thus robustly associated with the CE state at admission.

Proteins retained by the elastic-net model exhibited coordinated shifts in expression, with several demonstrating statistically significant differences after false discovery rate (FDR) correction (**Fig. 4**). These proteins spanned a range of effect sizes and directions, reflecting heterogeneous but structured alterations associated with CE rather than a small number of dominant univariate signals. Importantly, many proteins selected by elastic net would not have met stringent univariate significance thresholds, underscoring the ability of multivariate regularization to capture correlated, complementary signals characteristic of high-dimensional proteomic data.

**Figure 4.**
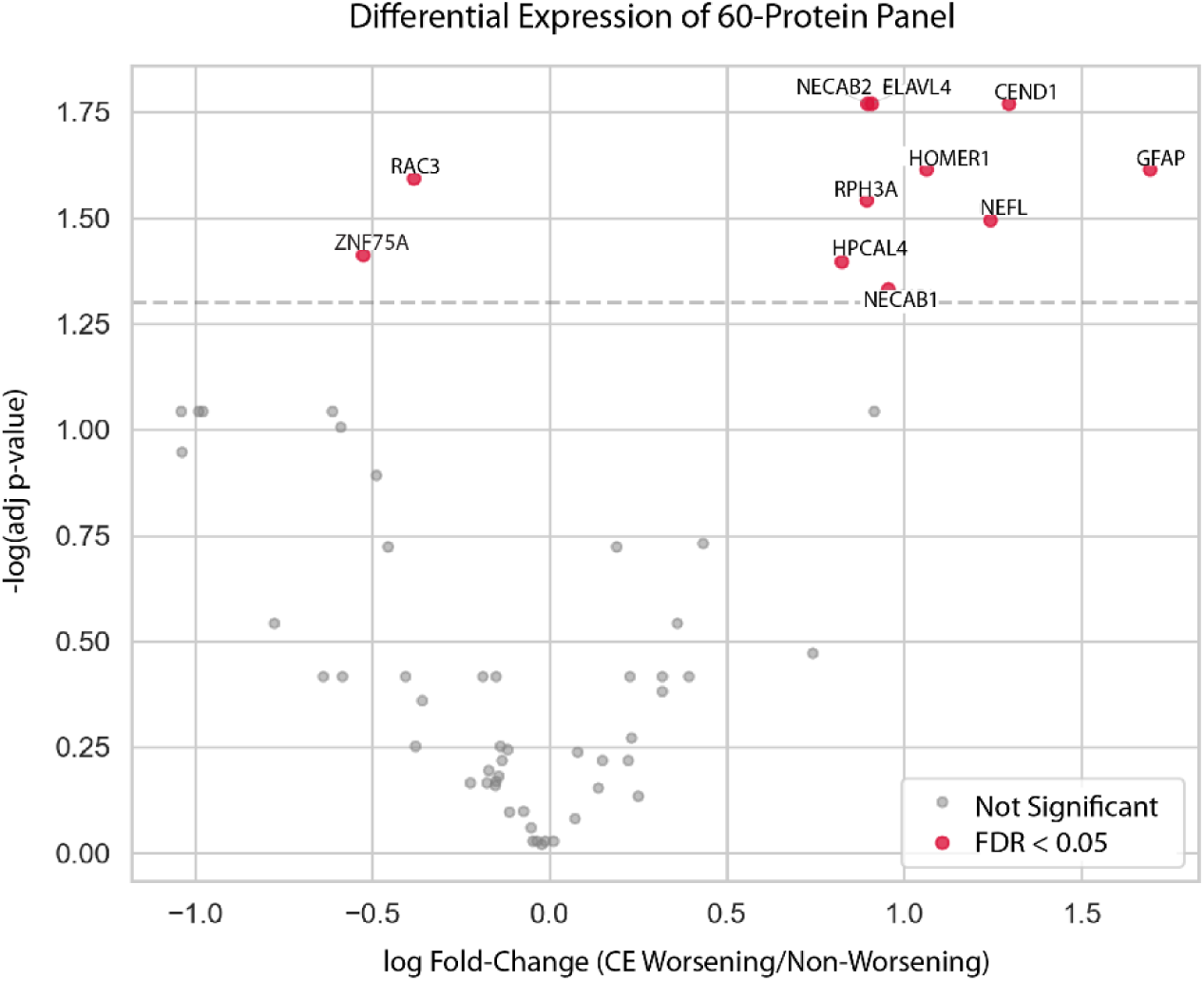
Differential expression of the 60-protein panel between patients with and without CE. Volcano plot showing log₂ fold-change in plasma protein expression between patients with and without CE plotted against the −log FDR-adjusted p-value. Proteins retained in the 60-protein panel by bootstrapped elastic-net regression are shown, with proteins meeting FDR significance (q < 0.05) highlighted in red and non-significant proteins shown in gray. Labeled proteins represent those with the largest effect sizes and/or statistical significance. Dashed horizontal line indicates the FDR significance threshold.

This expanded 60-protein panel was carried forward for unsupervised structure analysis, supervised prediction of subsequent CE worsening, and pathway enrichment analyses to support biological interpretation of CE-associated proteomic patterns.

### Baseline Proteomic Patterns Stratify Patients by CE Worsening

To evaluate the intrinsic structure of the expanded 60-protein panel identified by elastic-net regression, we performed PCA on baseline protein expression levels. PCA revealed a low-dimensional organization of the proteomic data, with the first two PCs accounting for 25.8% if the total variance across subjects (**Fig. 5 A; Supplementary Fig. 2 A**). When visualized in the PC1 x PC2 space, patients with CE on admission CT exhibited separation from those without CE, indicating that the multivariate proteomic signature distinguishes CE status at presentation beyond individual protein effects.

**Figure 5.**
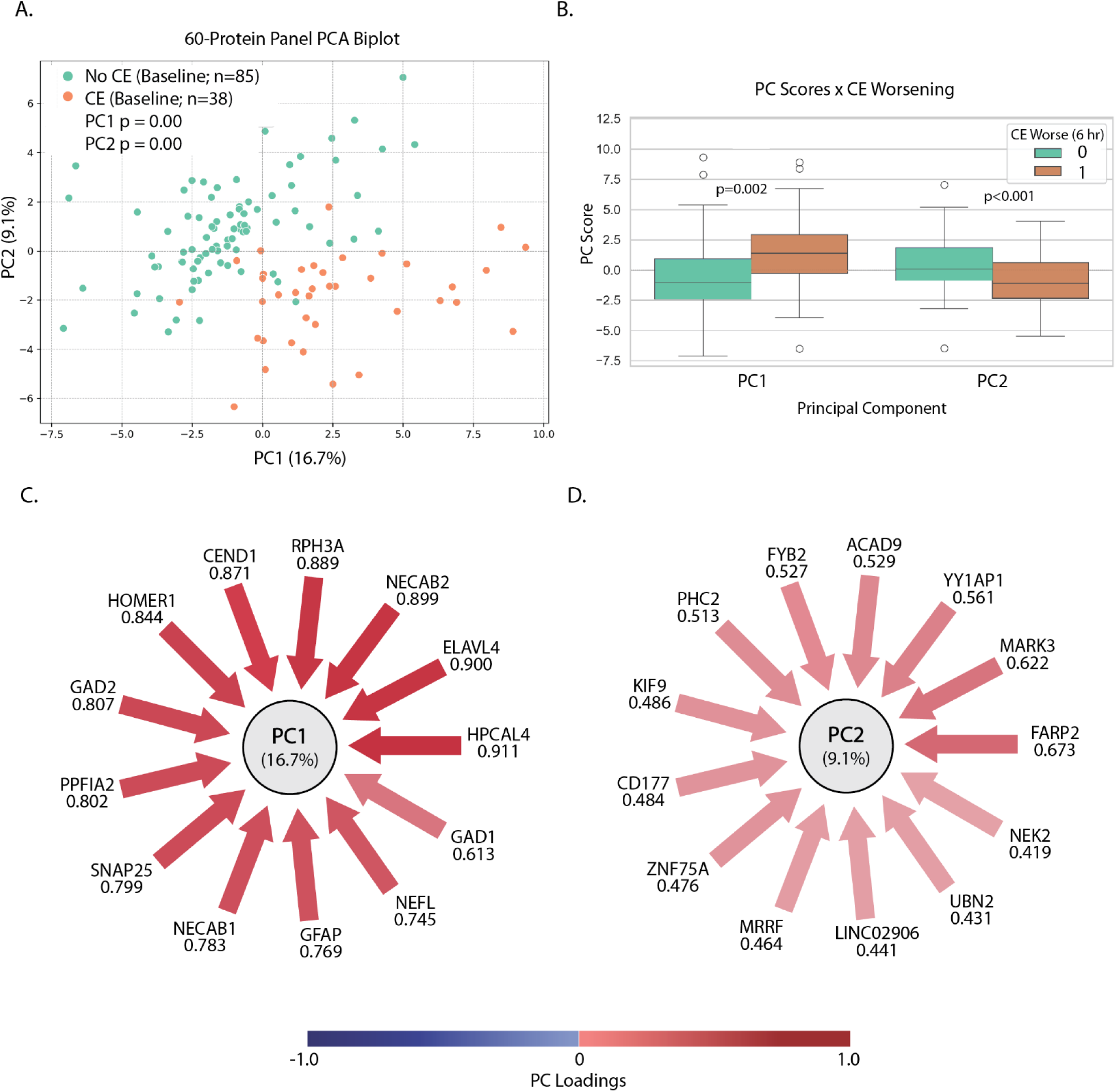
Baseline proteomic patterns associated with CE status and early CE worsening. (**A**) PCA of baseline expression levels for the elastic-net–derived 60-protein panel demonstrates partial separation between patients with and without CE on admission CT. (**B**) PC scores stratified by 6-hour CE worsening status show significantly higher PC1 scores and lower PC2 scores among patients who experienced CE worsening compared with those who did not (PC1: p = 0.002; PC2: p < 0.001). (**C**) Protein loadings contributing to PC1, which explains 16.7% of total variance. (**D**) Protein loadings contributing to PC2, explaining 9.1% of total variance.

We next assessed whether these latent proteomic dimensions were related to subsequent clinical trajectories. PC scores derived from baseline protein expression were compared between patients who did and did not experience radiographic CE worsening within 6 hours. Both PC1 and PC2 scores differed significantly between groups (PC1 p = 0.002; PC2 p < 0.001) demonstrating strong associations with subsequent CE worsening (**Fig. 5 B**) and suggesting that distinct axes of baseline proteomic variation are informative of early CE progression risk. Notably, PC1 scores were higher in the CE worsening group while PC2 scores were lower relative to the non-worsening group.

Inspection of PC loadings revealed biologically coherent protein contributions to each component (**Fig. 5 C,D**). PC1 (explaining 16.7% of total variance) was dominated by neuronal and synaptic proteins, including NECAB2, HPCAL4, ELAVL4, NEFL, SNAP25, and GAD1/2, as well as markers of astroglial activation such as GFAP (**Fig. 5 C; Supplementary Fig. 2 B**). Notably, PC1 scores were higher among patients who subsequently experienced CE worsening, indicating relative upregulation of this neuronal–glial injury–associated module at baseline. In contrast, PC2 (explaining 9.1% of variance) was characterized by a distinct set of proteins, including regulators of intracellular signaling, cytoskeletal organization, and cellular stress responses (**Fig. 5 D; Supplementary Fig. 2 C**), and PC2 scores were lower among patients who later worsened. Together, these findings indicate that the elastic-net–derived proteomic panel exhibits structured, low-dimensional organization at baseline, with multiple independent protein modules showing opposing directional associations with early CE worsening risk.

### Expanded Proteomic Panel Predicts Early Edema Worsening

To determine whether the expanded elastic-net–derived proteomic signature contained prognostic information independent of clinical variables, we trained supervised ML models using only the 60-protein panel as input features to predict radiographic CE worsening at 6 hours. Across classifiers, models trained on baseline protein expression demonstrated consistent discrimination between patients who did and did not experience worsening (**Fig. 6 A**).

**Figure 6.**
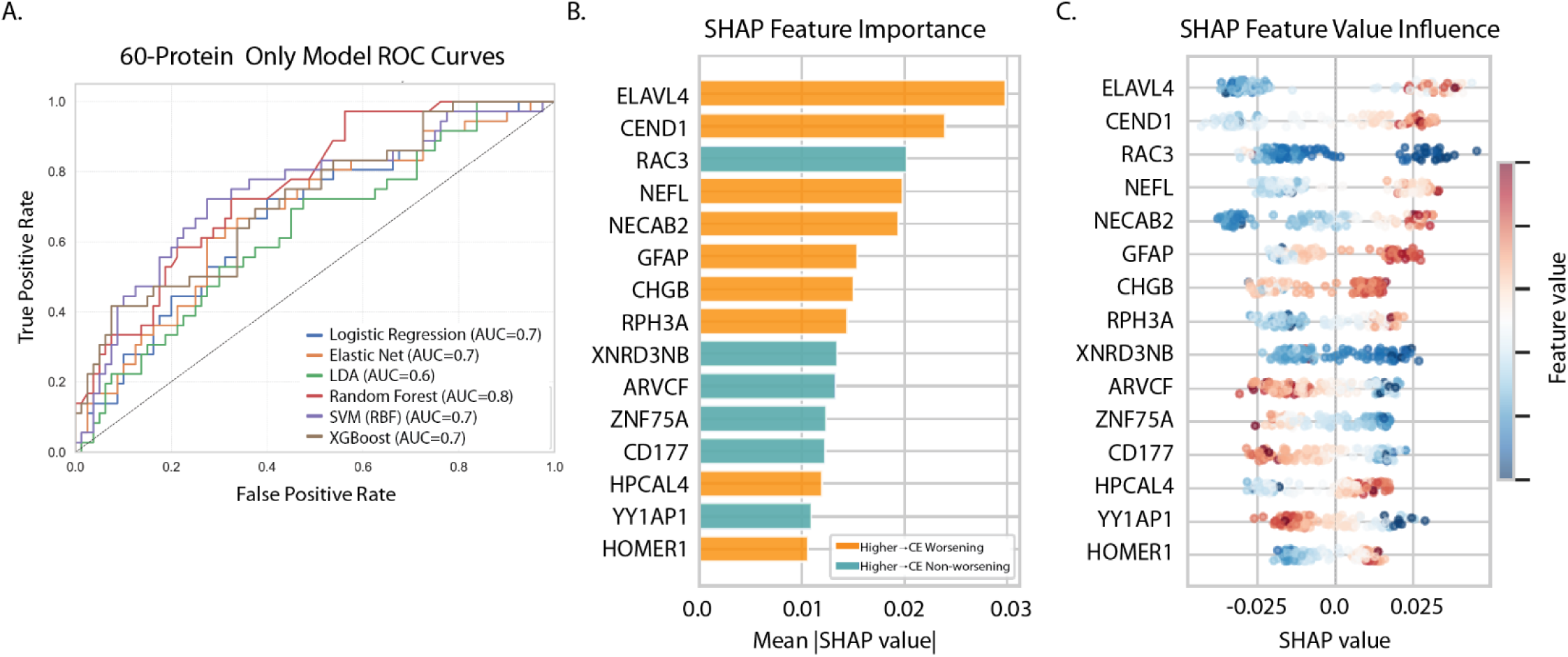
Prediction of early CE worsening using the 60-protein panel alone. **(A)**ROC curves comparing supervised ML classifiers trained using baseline expression of the elastic-net–derived 60-protein panel to predict radiographic CE worsening at 6 hours. Curves represent cross-validated model performance for logistic regression, elastic net, linear discriminant analysis (LDA), RF, SVM, and XGBoost. The RF classifier demonstrated the highest overall discrimination and was selected for downstream model interpretation and pathway enrichment analyses. The dashed diagonal line indicates chance-level performance. **(B).** RF feature attribution using SHAP for prediction of 6-hour CE worsening based on the 60-protein panel. Mean absolute SHAP values summarize the relative importance of top-ranked proteins, with color indicating the direction of association (red: higher expression associated with increased likelihood of CE worsening group; blue: higher expression associated with reduced likelihood of CE worsening group). **(C)** SHAP beeswarm feature importance plot showing the distribution of SHAP values across individuals for the top 15 protein contributors, colored by relative feature value (red: high expression; blue: low expression). Positive SHAP values indicate increased likelihood of CE worsening prediction, whereas negative SHAP values indicate decreased predicted likelihood of CE worsening.

Among the evaluated algorithms, the RF classifier achieved the highest overall discrimination (AUC = 0.76), with other models showing comparable but modestly lower performance. These findings indicate that baseline proteomic information alone captures clinically relevant signal associated with early CE worsening. Based on its superior discrimination and stability, the RF model was selected for downstream model interpretation and pathway-focused analyses in Goal 2.

### Interpretable Machine Learning Reveals Key Proteomic Patterns that Predict CE Worsening

To identify proteins most strongly contributing to model predictions of early CE worsening and to clarify the directionality of their effects, we interrogated the RF model using permutation-based feature importance and SHAP analysis. Both analyses revealed that individual proteins contributed to predicted CE worsening in opposing directions.

Higher expression of several neuronal, synaptic, and neuroglial injury–associated proteins, including ELAVL4, CEND1, NEFL, NECAB2, GFAP, and RPH3A, were associated with increased predicted risk of CE worsening (**Fig. 6 B,C; Supplementary Fig. 3**). In contrast, higher expression of a separate subset of proteins, including ARVCF, ZNF75A, CD177, RAC3, TXNRD3NB, and YY1AP1, were associated with lower predicted risk, indicating a protective or non-worsening–associated signature (**Fig. 6 B,C**). These opposing SHAP directions demonstrate that the expanded proteomic panel captures both CE risk-enhancing and risk-attenuating molecular signals at baseline. Furthermore, interaction terms within the SHAP analysis reveals that CE worsening may be driven by a broader network rather than strong, synergistic interaction across a small number of protein pairs.

Together, these findings indicate that early CE worsening is not driven by a single dominant biomarker but instead reflects the balance of multiple biological processes with opposing contributions to risk, reinforcing the value of multivariate proteomic modeling for mechanistic interpretation.

### Network analysis highlights coordinated neuronal–synaptic injury in patients with CE worsening

To contextualize the biological processes underlying proteins identified as high-impact contributors to early CE worsening, we performed network-based functional enrichment and pathway analysis using SHAP-prioritized proteins stratified by direction of association with outcome. Proteins associated with increased predicted risk of CE worsening formed a tightly interconnected network enriched for neuronal and synaptic signaling processes (**Fig 7**). Prominent pathways included chemical synaptic transmission, trans-synaptic signaling, synaptic vesicle docking and exocytosis, and gamma-aminobutyric acid (GABA) metabolic processes, with key hubs centered on synaptic and neurotransmission-related proteins such as GAD1, GAD2, SNAP25, HOMER1, and RPH3A.

**Figure 7.**
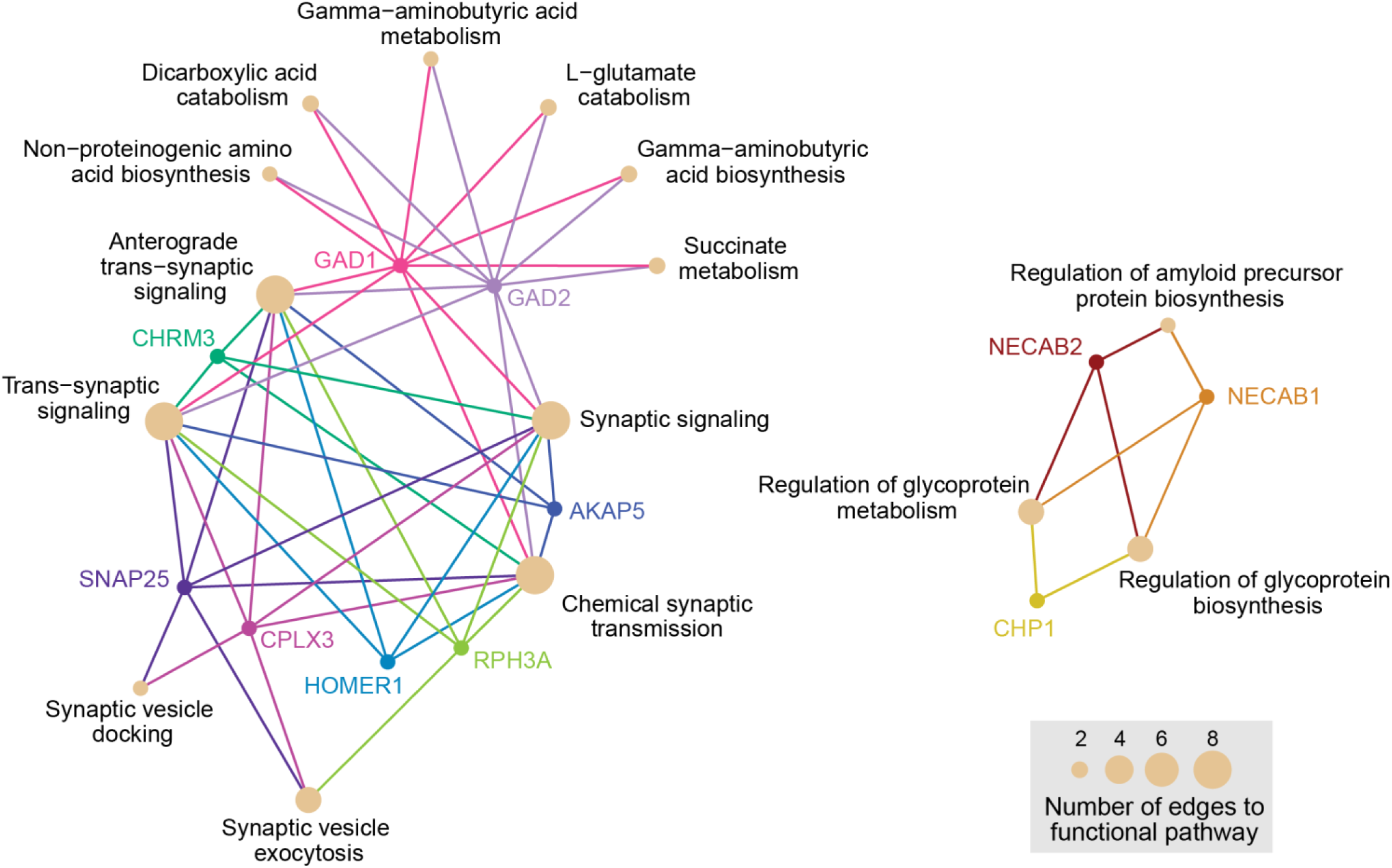
Functional network annotation analysis reveals coordinated neuronal–synaptic injury programs associated with CE worsening. Functional enrichment network constructed from SHAP-prioritized proteins associated with increased predicted risk of CE worsening. Nodes represent enriched biological processes or proteins, with edges indicating shared gene membership. The CE worsening-associated network is characterized by dense connectivity among neuronal and synaptic signaling pathways, including chemical synaptic transmission, trans-synaptic signaling, synaptic vesicle exocytosis, and GABA metabolic processes. Node size reflects pathway gene count. Colors correspond to individual proteins and their relevant pathways. Brown nodes and corresponding size is relative to the number of proteins assigned to membership in the listed pathway.

In contrast, proteins associated with reduced predicted risk of CE worsening did not form a comparably dense interaction network. Instead, these proteins were sparsely connected and mapped to more heterogeneous biological processes, including cellular signaling, metabolic regulation, and structural maintenance. This relative lack of network cohesion suggests that CE non-worsening trajectories may reflect preservation of cellular homeostasis rather than activation of a coordinated injury-specific molecular program.

Together, these findings indicate that early CE worsening is characterized by a coherent, multicomponent neuronal–synaptic injury network detectable at baseline, whereas CE non-worsening outcomes are associated with a more diffuse and less tightly organized proteomic landscape. This asymmetry supports a model in which early edema progression reflects convergence on specific neurobiological injury pathways rather than the absence of injury signals.

## Discussion

The heterogeneity of TBI has stymied the field of neurotrauma for decades, contributing to disappointing clinical trial results and lack of preventative treatments for CE. However, with the advent of blood-based biomarkers in use for patient stratification for imaging clinically, and for entry into clinical treatment trials, the potential for new sensitive and specific blood-based biomarkers is compelling and timely.^28–30^ In this study, we leveraged high throughput multiplex immunoassay proteomics and ML to pursue two complementary goals in a prospective cohort of patients with low GCS TBI and intracranial hemorrhage: (1) to identify a set of candidate biomarkers and develop a predictive model that could be used to assess CE risk at admission for critically ill patients after TBI and (2) to conduct a mechanistic, hypothesis-generating investigation to identify the molecular networks involved in CE progression in the first hours post-injury.

Our efforts fit readily into the new characterization of TBI (CBI-M), to advance the biomarker pillar.^31^ The current biomarker landscape in TBI has been largely shaped by the development and FDA authorization of GFAP and UCH-L1, milestones that have demonstrated the feasibility of measuring circulating proteins for guidance in acute clinical decision making.^28,32^ However, those assays are currently limited to predicting the absence of intracranial lesions on CT in mild TBI for the purposes of binary triage decisions. Our work substantially extends this paradigm in several ways. First, we target a distinct and clinically urgent question: can circulating protein signatures detectable upon admission predict worsening CE over ensuing hours. Second, rather than relying on established neurotrauma markers, we demonstrate the feasibility of employing an unbiased, proteome-wide screen of over 5400 proteins to discover novel, combinatorial signatures associated with evolving pathologies following moderate-severe TBI. This approach resulted in protein panels including several markers not previously associated with CE pathology such as ELAVL4, CEND1, NECAB2, HPCAL4, and RPH3A alongside established neurotrauma markers such as GFAP and NEFL (also known as NFL) and known synaptic markers such as SNAP25.^33–36^ The majority of proteins identified were elevated in the CE-worsening group which is consistent with many blood-based biomarkers used to quantify injury burden following TBI.^33,34,37^ Interestingly, emerging evidence suggests CEND1 expression is downregulated in brain tissue following experimental TBI, and its deficiency drives presynaptic mitochondrial dysfunction and cognitive impairments in models of Alzheimer’s Disease.^38,39^ The peripheral elevation observed here may reflect injury-dependent release from damaged neurons, though this remains to be confirmed.

In designing our predictive modeling framework in the first phase of the current work, we attempted to maximize discrimination and limit the incidence of false negatives while minimizing the number of proteins required to achieve optimal performance to support a feasible, parsimonious protein panel that could be brought forward in the future for clinical translation. We felt that it was important to develop a panel that could be sent shortly after injury, at the emergency department admission timepoint to facilitate triage and patient stratification, as this may enable earlier identification of patients at risk for CE worsening than imaging or clinical variables alone. The 12-protein panel, in this case, represents a feasible target for focused assay development and prospective validation, particularly as rapid multiplex immunoassay platforms continue to advance toward point-of-care development. When combined with six routinely available clinical admission variables and optimized to avoid false negatives (i.e., categorizing a patient as a non-worsener when they will experience CE worsening), this panel yielded a model with good discrimination (AUC = 0.78) and high sensitivity (recall = 0.83) for predicting radiographic CE at 6 hours. Importantly, this combination of protein and clinical features outperformed models trained using either the clinical or proteome information alone reinforcing the idea that molecular and clinical data provide complementary, non-redundant, information for CE risk prediction. Moreover, the 12-protein + clinical model achieved a sensitivity of 83% alongside a specificity of 68% (XGBoost) at the optimized threshold, with alternative classifiers yielding a more balanced profile (LDA: 71% sensitivity, 74% specificity), discriminative profiles substantially more balanced than currently deployed single- and dual-analyte approaches, which, while highly sensitive (>95%), are constrained by specificity as low as 36%.^29^ This advantage is consistent with evidence that multi-analyte protein panels often outperform individual markers for TBI outcome prediction.^40,41^

While CE takes hours to days to come to full expression, the pathophysiology begins early after injury. Recently, a group of investigators demonstrated that injury severity as measured by the injury severity score could predict the time of onset of CE. In patients with mild/moderate TBI (AIS ≤ 3), the probability of CE occurring at 25%, 50%, and 75% was found to be at 12.8 h, 19.7 h, and 34.5 h from injury, respectively while patients with severe TBI (AIS > 3), the probabilities of occurrence were earlier, at 8.1 h, 12.5 h, and 21.6 h for 25%, 50%, and 75% probabilities.^42^ Currently, no validated blood-based biomarkers exist to anticipate CE worsening or to illuminate the molecular processes driving divergent edema trajectories in individual patients. Our findings suggest that early CE is associated with a coordinated neuronal–synaptic injury signature detectable at admission. This observation is in keeping with the literature with brain biopsy tissue post-severe TBI showing degenerative synaptic changes, and recent studies of CSF revealing elevated levels of synaptic proteins in CSF post-TBI in critically ill patients with extraventricular drains.^43,44^ We extend these findings to circulating plasma at the hyperacute, admission timepoint where we similarly observed proteins associated with CE-worsening converge on pathways related to synaptic transmission, vesicle cycling, calcium signaling, and axonal integrity such as SNAP25. Notably, NfL, has emerged as a robust circulating marker of neuroaxonal injury across multiple neurodegenerative conditions and has shown prognostic utility in TBI, correlating with injury severity, brain atrophy, and long-term outcomes.^34,35^ Circulating NfL has now been validated as a non-specific but highly sensitive marker of neuroaxonal damage across the full spectrum of TBI severity, with levels peaking at 10 days to 6 weeks post-injury and remaining elevated at one year in moderate-severe TBI.^35^ Its inclusion in the FDA’s Breakthrough Device Designation pathway for neurodegenerative disease monitoring further underscores its translational maturity as a neuronal injury biomarker.^45,46^ Our findings broaden the relevance of NfL from a general marker of axonal integrity to a component of a multi-protein signature specifically associated with early CE progression. Further, the co-elevation of astroglial (GFAP) and neuronal (NfL, SNAP25) markers in the CE-worsening group is consistent with emerging models of neurovascular unit disruption in early TBI, where astrocytic swelling and synaptic dysfunction co-occur as initiating events in cytotoxic edema formation.^6,7^

Viewed in aggregate, the 60-protein panel is notable for how strongly it localizes to biological systems expected to be relevant in the earliest stages of secondary injury-related pathophysiological cascades and edema progression: synaptic excitability, calcium-sensitive signaling, vesicle trafficking, and astroglial responses.^6–8,47–50^ Rather than pointing primarily to generalized tissue destruction, the CE worsening-associated signature is enriched for proteins involved in glutamatergic and GABAergic signaling, synaptic vesicle dynamics, and calcium-dependent neuronal stress responses. Several high-impact proteins illustrate this biology. ELAVL4 (HuD) is a neuronal RNA-binding protein with canonical roles in neuronal development and synaptic plasticity.^51^ ELAVL4/HuD has been implicated in multiple neurodegenerative conditions, including Parkinson’s, Alzheimer’s, and amyotrophic lateral sclerosis, where altered ELAVL4/HuD regulation disrupts target transcripts involved in synaptic function.^51,52^ Further, a growing body of evidence highlights HuD as a key mediator of neuronal plasticity, particularly in the context of axonal repair.^51–54^ HOMER1 is a postsynaptic scaffold linking metabotropic glutamate receptors to intracellular calcium signaling pathways. Notably, HOMER1 has been shown to be neuroprotective following experimental TBI by attenuating mGluR1-mediated excitotoxicity.^55^ Its expression is induced within hours of experimental TBI, likely in proportion to the severity of excitotoxic challenge.^55^ The elevated circulating levels observed in the CE-worsening group may reflect the intensity of the underlying excitotoxic insult rather than a deleterious effect of the protein itself. Proteins involved in synaptic vesicle trafficking and neurotransmitter release, including SNAP25, CPLX3, and RPH3A, further localize the signal to presynaptic machinery that is tightly coupled to calcium influx and excitotoxic stress.^56–59^ Complementing these synaptic proteins, neuronal calcium-binding proteins such as NECAB2 and HPCAL4 suggest perturbation of intracellular calcium regulation, a central feature of early cytotoxic injury cascades.^60,61^ Together with astroglial markers such as GFAP, these findings suggest that patients who worsen may be characterized by a hyperacute proteomic state marked by coupled synaptic dysfunction, calcium disequilibrium, and astroglial participation in the earliest phases of edema evolution. Importantly, although many of these proteins have not previously been linked directly to post-TBI CE, their established roles in synaptic transmission, calcium-dependent signaling, receptor trafficking, excitotoxic injury, and astrocyte-associated injury biology make them plausible components of the hyperacute molecular environment in which cytotoxic and ionic edema emerge.

In contrast, proteins associated with reduced predicted risk of CE worsening, including ARVCF, ZNF75A, CD177, AC3, TXNRD3NB, and YY1AP1, did not form a comparably dense interaction network in the functional enrichment analyses performed. Instead, these proteins mapped to heterogenous biological processes including cellular signaling, metabolic regulation, and structural maintenance. Notably, CD177, a glycosylphosphatidylinositol-anchored glycoprotein expressed on a subset of neutrophils, was recently identified as a marker of an anti-inflammatory neutrophil population that is neuroprotective following CNS injury, including TBI, where adoptive transfer of CD177+ neutrophils attenuated neuroinflammation.^62^ Its association with reduced CE risk in our data may reflect a peripheral immune phenotype that limits secondary inflammatory injury and edema progression. The relative absence of a dense network in the non-worsening group may suggest preservation of homeostasis rather than the activation of a coordinated protective program. This asymmetry reinforces the interpretation that early CE worsening reflects convergence on specific neurobiological injury pathways, while CE stability may reflect the absence of coordinated disruption.

In the context of acute neurotrauma, integrating ML with structured biological interpretation offers a way to extract mechanistic insight from predictive models, rather than treating model performance as an end in itself. In fact, a deliberate design feature of this study was the separation of clinically feasibly prediction (Goal 1) from hypothesis-generating biological exploration (Goal 2) within a unified analytical framework. Clinical translation demands parsimony, interpretability, and feasibility, whereas mechanistic discovery benefits from broader feature space exploration and network-level interrogation. This dual-analytic strategy enabled us to avoid the common tension in biomarker studies between overfitting a discovery panel to a clinical endpoint and sacrificing biological depth for the sake of model simplicity. ML algorithms, particularly methods like those best performing in this study (XGBoost and Random Forest), have demonstrated advantages over traditional regression approaches for outcome prediction in biomedical applications, largely due to their capacity to capture non-linear relationships, feature interactions, and high-dimensional data structures.^63,64^ Importantly, we employed a rigorous nested cross-validation framework to minimize information leakage and optimism bias.^20,65,66^ Further, the integration of SHAP-based model interpretation with structured biological annotation transforms what can otherwise be opaque predictive models into sources of mechanistic insight. By quantifying the direction and magnitude of protein contribution to individual predictions and mapping high-impact features onto function interaction networks, we were able to bridge the gap between model performance and biological understanding. Adherence to reporting standards for prediction models, including the TRIPOD statement, informed the presentation of model performance and internal validation in this work This approach exemplifies how integrating ML with structured biological methods can yield both clinically actionable tools and hypothesis generating mechanistic insight in acute neurotrauma.^20^

Several limitations warrant consideration. This investigation was focused on a hyperacute time point early after the initial injury. We do not have the benefit of serial blood sampling to follow the trajectory of the candidate protein biomarkers over time, nor do we have serial imaging to follow the evolution of CE. Future investigations would benefit from protocolized imaging with synchronized blood sampling to follow the natural history of CE, which peaks much later than the first 6 hours that this investigation monitors. Our binary definitions of CE and worsening CE have the potential for misclassification in that our cut-offs may have missed clinically significant CE that was just under the threshold for midline shift or basal cistern compression. Future studies may incorporate deep learning techniques in two complementary ways: first, as an additional input, where imaging-derived volumetric features extracted from an admission scan are combined with the circulating proteomic signature to create a more informative, multimodal predictor of CE trajectory; and second, to more accurately and reproducibly measure CE on imaging replacing the binary classification of CE worsening used in the present study.^67–69^ Additionally, all patients included in this investigation were enrolled at a single center in Portland, Oregon USA. While the demographics reflect TBI (predominantly male sex), and the racial and ethnic composition of the site of enrollment, these qualities may also impact generalizability. Further, ML classifiers generally benefit from large training samples, and the modest sample size in this study limits statistical precision, as reflected in the moderate confidence intervals around AUC estimates, and increases susceptibility to overfitting. All performance metrics reported are based on internal cross-validation. Future external validation in an independent TBI cohort is needed to confirm generalizability prior to prospective clinical development.

By applying interpretable ML to admission-time plasma proteomics, our results suggest that early CE progression reflects coordinated disruption of neuronal–synaptic signaling, calcium-dependent stress responses, and astroglial injury processes, a molecular signature detectable in peripheral blood within hours of trauma. Conceptually, these results support a shift from viewing circulating biomarkers solely as indicators of injury burden toward using multidimensional molecular signatures to characterize evolving biological states after TBI. If validated in independent cohorts, such approaches could enable biology-informed risk stratification that could identify patients at heightened risk for early edema progression, guided targeted enrollment in prevention trials, and providing new insight into the molecular pathways amenable to therapeutic intervention.

## Conclusion

In this hypothesis generating investigation, we have proposed candidate biomarkers which may provide clinical decision support and/or mechanistic inight into the formation and treatment of cerebral edema in critically ill patients after TBI, if our findings can be validated in an independent cohort of patients and followed by rigorous implementation testing. This work has the potential to improve the integration of clinical, biomarkers, and imaging data to support multimodal clinical decision making.

## Data Availability

Data sharing upon reasonable request to corresponding author. All code for data processing and analyses is available at https://github.com/Hinson-Lab/Cerebral-Edema. The blood-based biomarker data processing pipeline and statistical analyses are implemented in Python and R with detailed documentation for reproducibility. Interactive Enrichment Analysis developed by the Gladstone Bioinformatics Core can be accessed at https://github.com/gladstone-institutes/Interactive-Enrichment-Analysis.

## Acknowledgements

We thank the Gladstone Bioinformatics Core for their interactive enrichment analysis tools.

