## Supplementary Figures 1-4 for "Protein Predictors of Worsening Cerebral Edema in Traumatic Brain Injury in Critically Ill Patients"

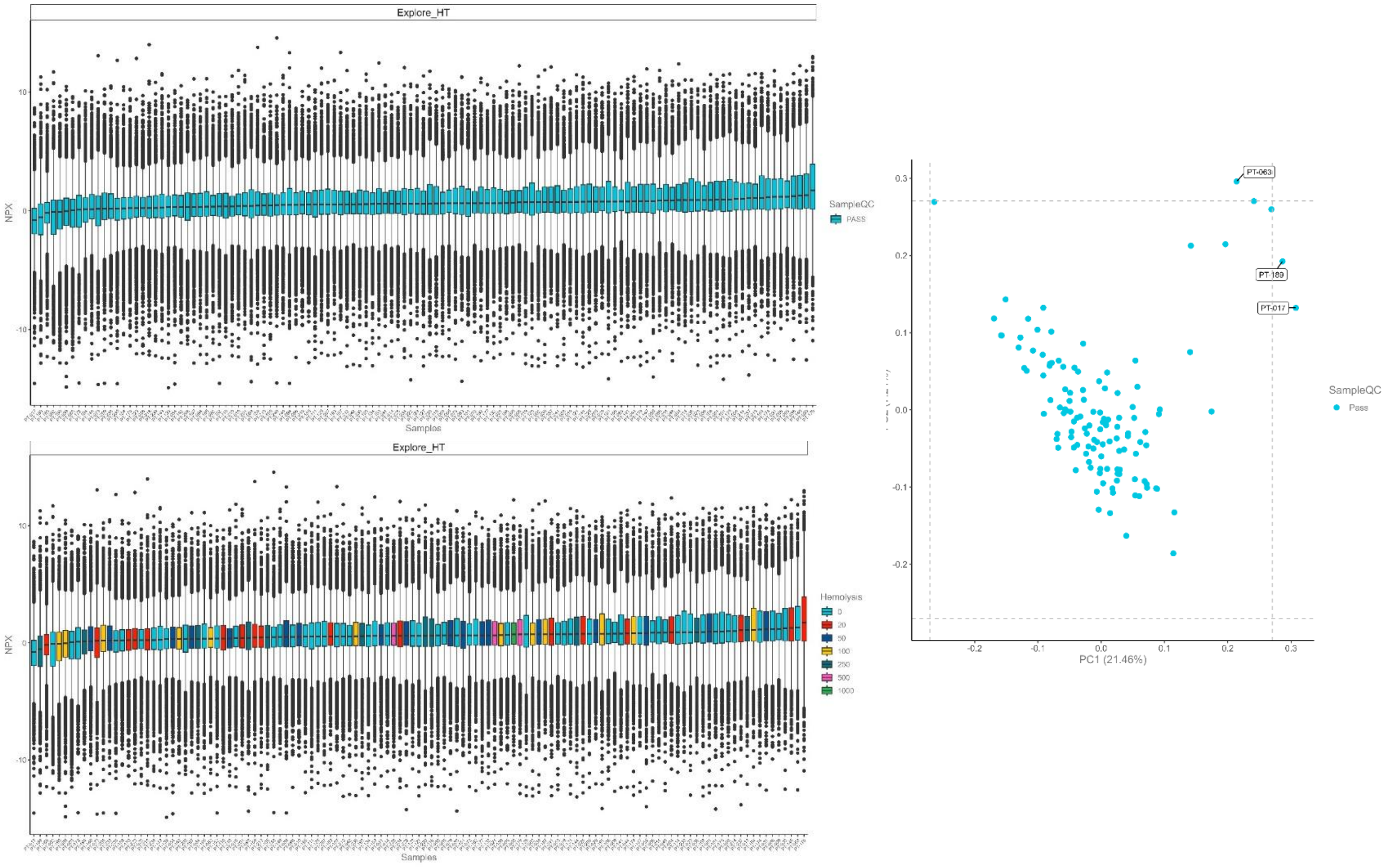


**Supplementary Figure 1. OLink PEA QC plots.** (A) NPX values across samples (B) stratified by hemolysis status. (C) PCA investigating global structure for evidence of technical outliers.

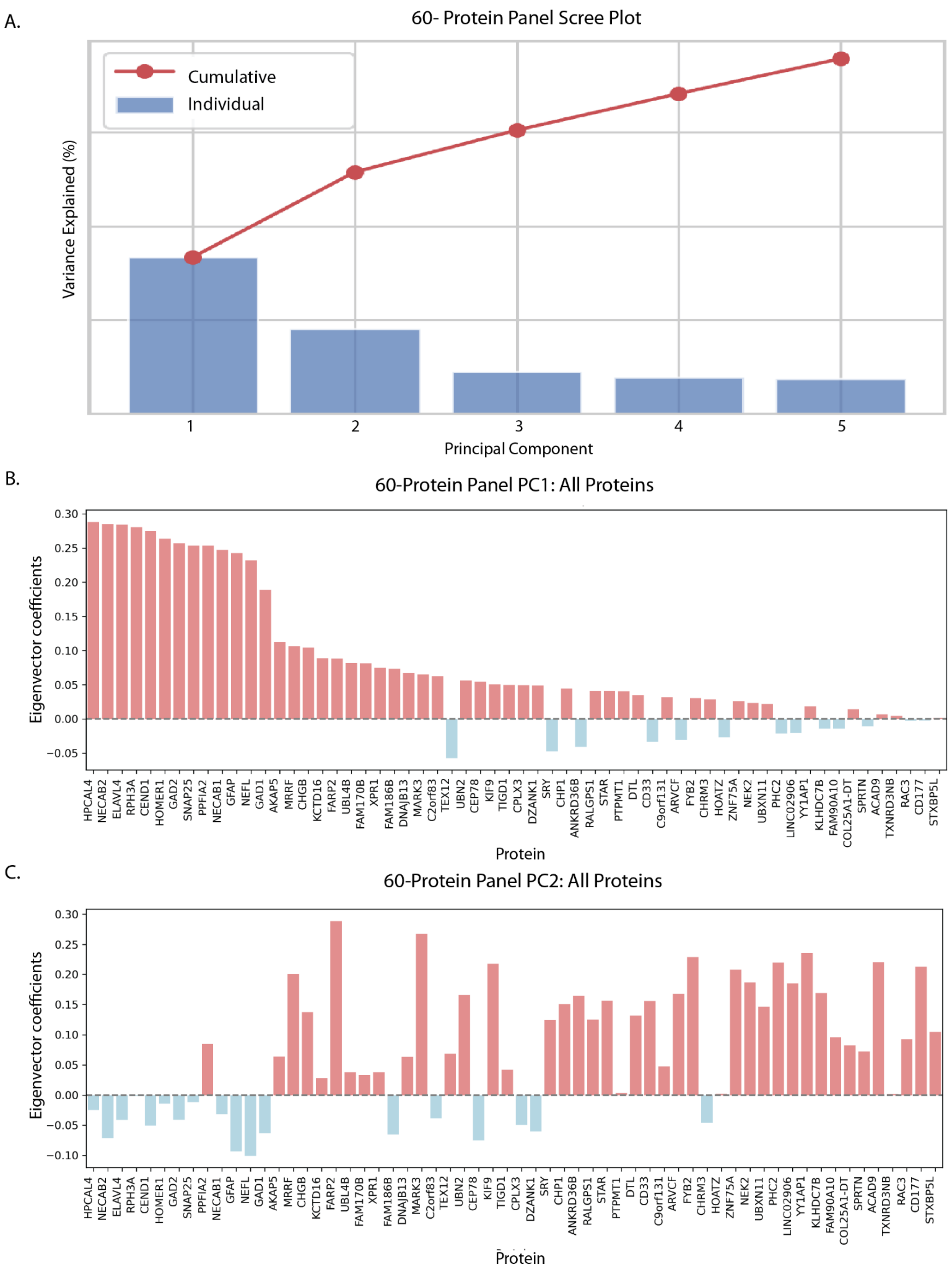


**Supplemental Fig 2. PCA characteristics for 60-protein mechanistic panel.** (A) Scree plot visualizing amount of variance explained by each PC. (B) Eigenvector coefficients describing each protein's contribution to PC1. (C) Eigenvector coefficients describing each protein's contribution to PC2.

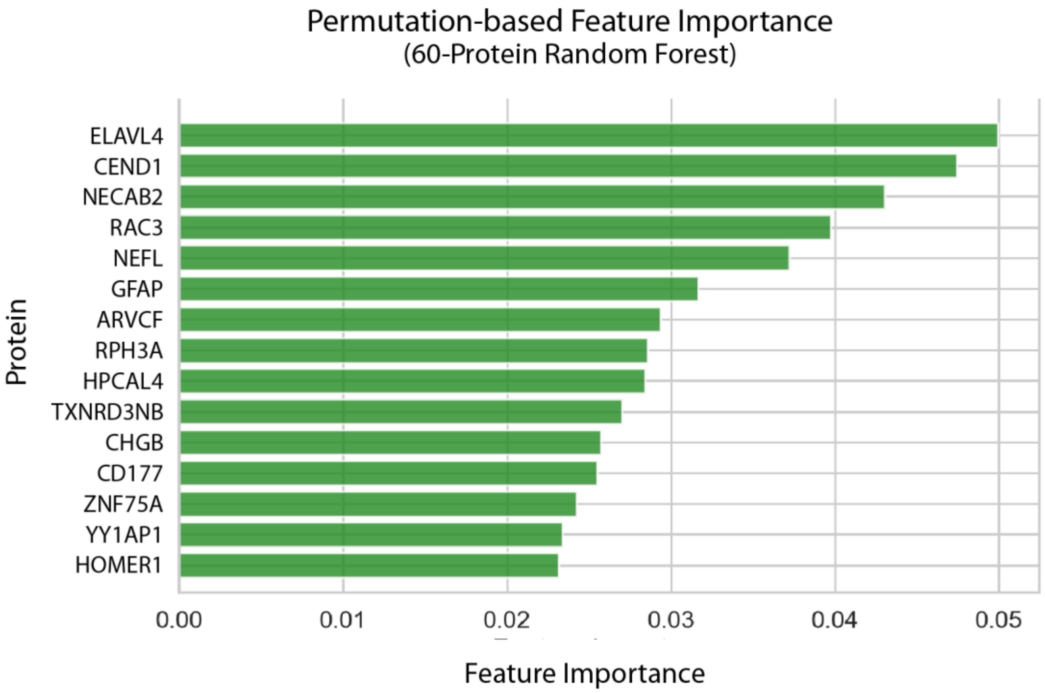


**Supplemental Fig 3. Random Forest permutation based feature importance.** Relative importance corroborates SHAP values emphasizing substantial discriminatory ability of subset of proteins for detecting CE worseners from non-worseners.

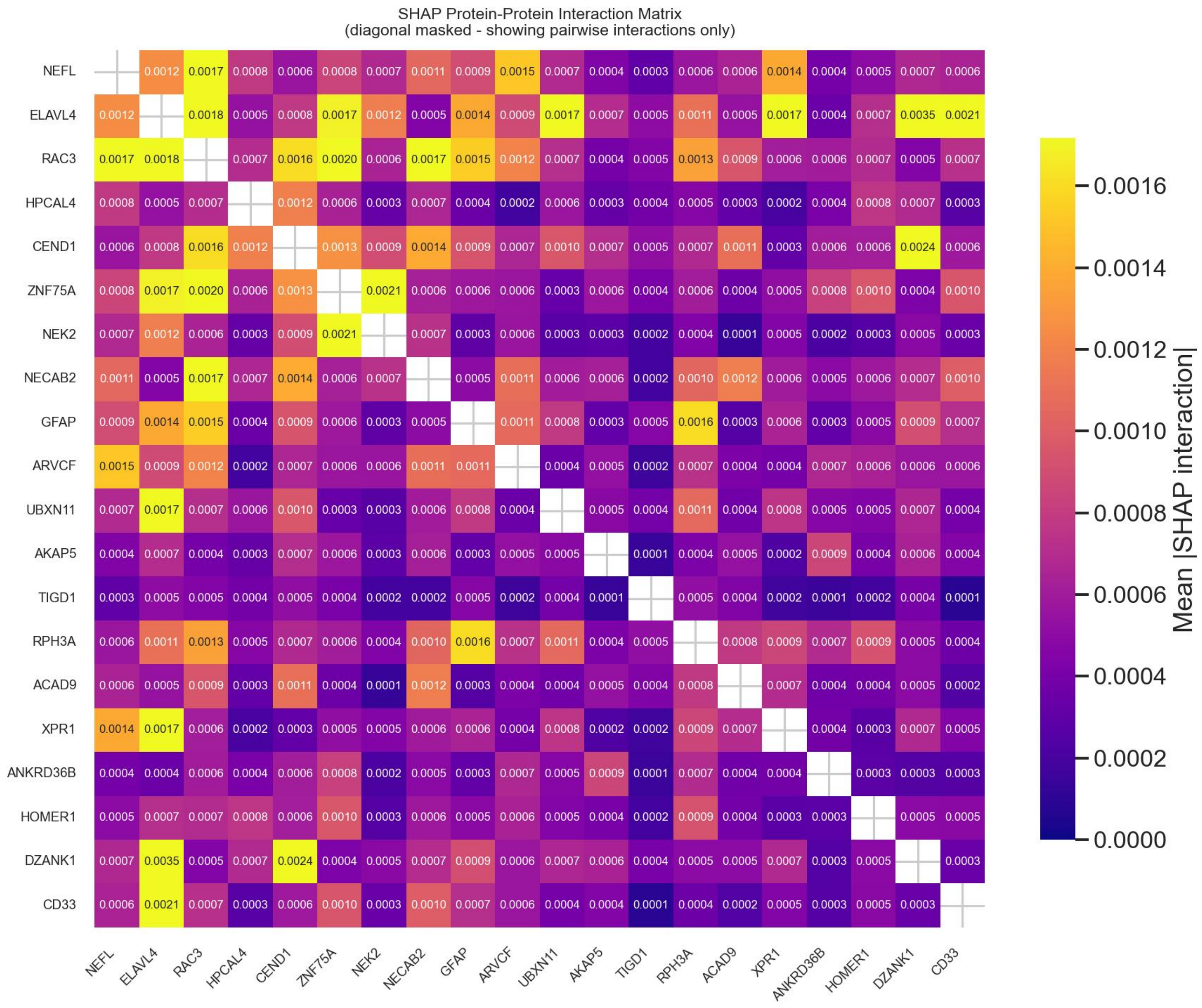


**Supplementary Figure 4.** SHAP interaction heatmap. Lack of large interaction terms suggests that CE worsening is not driven by a few dominant synergistic protein pairs
